# Inferring Phylogenetic Networks Using PhyloNet

**DOI:** 10.1101/238071

**Authors:** Dingqiao Wen, Yun Yu, Jiafan Zhu, Luay Nakhleh

## Abstract

PhyloNet was released in 2008 as a software package for representing and analyzing phylogenetic networks. At the time of its release, the main functionalities in PhyloNet consisted of measures for comparing network topologies and a single heuristic for reconciling gene trees with a species tree. Since then, PhyloNet has grown significantly. The software package now includes a wide array of methods for inferring phylogenetic networks from data sets of unlinked loci while accounting for both reticulation (e.g., hybridization) and incomplete lineage sorting. In particular, PhyloNet now allows for maximum parsimony, maximum likelihood, and Bayesian inference of phylogenetic networks from gene tree estimates. Furthermore, Bayesian inference directly from sequence data (sequence alignments or bi-allelic markers) is implemented. Maximum parsimony is based on an extension of the “minimizing deep coalescences” criterion to phylogenetic networks, whereas maximum likelihood and Bayesian inference are based on the multispecies network coalescent. All methods allow for multiple individuals per species. As computing the likelihood of a phylogenetic network is computationally hard, PhyloNet allows for evaluation and inference of networks using a pseudo-likelihood measure. PhyloNet summarizes the results of the various analyses, and generates phylogenetic networks in the extended Newick format that is readily viewable by existing visualization software, [phylogenetic networks; reticulation; incomplete lineage sorting; multispecies network coalescent; Bayesian inference; maximum likelihood; maximum parsimony.]

---

With the increasing availability of whole-genome and multi-locus data sets, an explosion in the development of methods for species tree inference from such data ensued. In particular, the multispecies coalescent (Degnan and Rosenberg, 2009) played a central role in explaining and modeling the phenomenon of gene tree incongruence due to incomplete lineage sorting (ILS), as well as in devising computational methods for species tree inference in the presence of ILS; e.g., (Heled and Drummond, 2010; Liu, 2008).

Nevertheless, with the increasing recognition that the evolutionary histories of several groups of closely related species are reticulate (Mallet et al., 2016), there is need for developing methods that infer species phylogenies while accounting not only for ILS but also for processes such as hybridization. Such reticulate species phylogenies are modeled by *phylogenetic networks* (Nakhleh, 2010). A phylogenetic network extends the phylogenetic tree model by allowing for horizontal edges that capture the inheritance of genetic material through gene flow (Fig. 1(a)). While the phylogenetic network captures how the species, or populations, have evolved, gene trees growing within its branches capture the evolutionary histories of individual, recombination-free loci (Fig. 1(b)). The relationship between phylogenetic networks and trees is complex in the presence of ILS (Zhu et al., 2016). Mathematically, the topology of a phylogenetic network takes the form of a rooted, directed, acyclic graph. In particular, while gene flow involves contemporaneous species or populations, past extinctions or incomplete sampling for taxa sometimes result in horizontal edges that appear to be “forward in time” (Fig. 1). It is important to account for such an event, which is why acyclicity, rather than having truly horizontal edges, is the only constraint that should be imposed on rooted directed graphs, in practice, if one is to model reticulate evolutionary histories.

**FIGURE 1.**
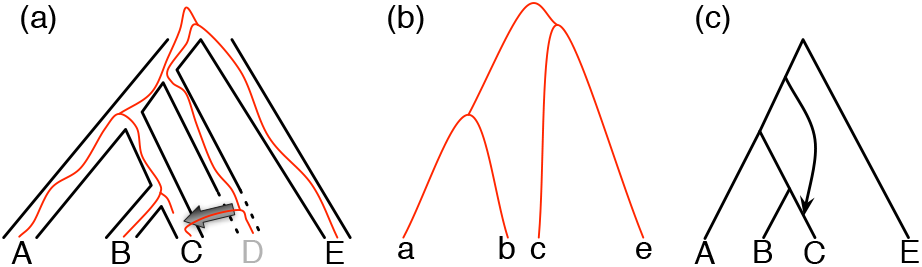
(a) A phylogenetic network on fix taxa, with taxon *D* missing (due to extinction or incomplete sampling), and a hybridization involving (ancestors of) taxa *D* and *C*. Shown within the branches of the phylogenetic network is a tree of a recombination-free locus whose evolutionary history includes introgression. (b) The gene tree that would be estimated, barring inference error, on the locus illustrated in (a), (c) An abstract depiction of the phylogenetic network of (a) given that taxon *D* is missing.

For inference of phylogenetic networks from multilocus data sets, the notions of coalescent histories and the multispecies coalescent were extended to phylogenetic networks (Yu et al., 2012, 2011). Based on these new models, the “minimizing deep coalescence” criterion (Maddison, 1997; Than and Nakhleh, 2009) was extended to phylogenetic networks, which allowed for a maximum parsimony inference of phylogenetic networks from the gene tree estimates of unlinked loci (Yu et al., 2013a). Subsequently, maximum likelihood inference (from gene tree estimates) via hill-climing heuristics and Bayesian inference via reversible-jump Markov chain Monte Carlo (RJMCMC) were devised (Wen et al., 2016; Yu et al., 2014). As computing the likelihood of a phylogenetic network formed a major bottleneck in the inference, speedup techniques for likelihood calculations and pseudo-likelihood of phylogenetic networks were introduced (Yu and Nakhleh, 2015b; Yu et al., 2013b). Finally, to enable direct estimation from sequence data, new methods were developed for Bayesian inference from sequence alignments of unlinked loci (Wen and Nakhleh, 2017) as well as bi-allelic markers of unlinkedloci (Zhu et al., 2017). Here we introduce PhyloNet 3, a software package for phylogenetic network inference from multi-locus data under the aforementioned models and criteria. This version is a significant expansion of the version reported on in (Than et al., 2008). Phylogenetic networks inferred by PhyloNet are represented using an extended Newick format and can be readily visualized by Dendroscope (Huson and Scornavacca, 2012).

## MODELS AND MAIN INFERENCE FEATURES

### Simple Counts of Extra Lineages: Maximum Parsimony

Minimizing the number of deep coalescences, or MDC, is a criterion that was proposed originally by Maddison (Maddison, 1997) for species tree inference and later implemented and tested in both heuristic form (Maddison and Knowles, 2006) and exact algorithms (Than and Nakhleh, 2009). Yu *et al*. (Yu et al., 2013a) extended the MDC criterion to phylogenetic networks. The **InferNetwork_MP** command infers a species network with a specified number of reticulation nodes under the extended MDC criterion. Inference under this criterion is done via a local search heuristic, and the phylogenetic networks returned by the program include, in addition to the topologies, the inheritance probability estimates, as well as the number of extra lineages on each branch of the network, and the total number of extra lineages of the phylogenetic network. For this program, only gene tree topologies are used as input (that is, gene tree branch lengths are irrelevant), and the number of individuals per species could vary across loci. Furthermore, to account for uncertainty in the input gene tree estimates, the program allows for a set of gene trees per locus that could be obtained from a bootstrap analysis or a posterior sample on the sequences of the respective locus. For inference under the MDC criterion, the maximum number of reticulation events in the phylogenetic network must be specified *a priori*. Full details of the MDC criterion for phylogenetic networks and the inference heuristics can be found in (Yu et al., 2013a).

Inference based on the MDC criterion does not allow for estimating branch lengths or any other associated parameters of the inferred phylogenetic network beyond the topology and inheritance probabilities.

### When the Species Phylogeny’s Branches Are Too Short: Maximum Likelihood

One limitation of inference based on the MDC criterion is the inability to estimate parameter values beyond the network’s topology. Another limitation is the fact that such inference is not statistically consistent for species trees (Than and Rosenberg, 2011), which implies problems in the case of phylogenetic network inference based on the criterion as well (more on the notion of “statistical consistency” in the case of networks below). The latter problem arises especially when the species phylogeny has very short branches. To address these two limitations, Yu *et al*. (Yu et al., 2014) implemented maximum likelihood estimation of phylogenetic networks based on the multispecies network coalescent (Yu et al., 2012). The **InferNetwork_ML** command infers a maximum likelihood species network(s) along with its branch lengths (in coalescent units) and inheritance probabilities. During the search, the branch lengths and inheritance probabilities of a proposed species network can be either sampled or optimized (the former is much faster and has been shown to perform very well). The input consists of either rooted gene tree topologies alone, or rooted gene trees with branch lengths (in coalescent units). If the gene tree branch lengths are to be used, the gene trees must be ultrametric. As in the case of maximum parsimony inference, local search heuristics are used to obtain the maximum likelihood estimates. Furthermore, multiple individuals per species could be used, and their numbers could vary across loci. Multiple gene trees per locus could be used, as above, to account for uncertainty in the gene tree estimates. The user can either specify the maximum number of reticulation events *a priori* or utilize the cross-validation (the **InferNetwork_ML_CV** command) or bootstrap (the **InferNetwork_ML_Bootstrap** command) to determine the model complexity. Furthermore, several information criteria (AIC, BIC, and AICs) are implemented. Full details of the maximum likelihood inference of phylogenetic networks and the inference heuristics can be found in (Yu et al., 2014).

It is important to note that computing the likelihood of a phylogenetic network is a major computational bottleneck in all statistical inference methods implemented in PhyloNet. To ameliorate this problem, PhyloNet also allows for inference of phylogenetic networks based on a “pseudo-likelihood” measure, via the **InferNetwork_MPL** command. However, for this method, the input could consist only of gene tree topologies (branch lengths are not allowed). Multiple individuals per species, as well as nonbinary gene trees, are also allowed. Full details about inference under pseudo-likelihood can be found in (Yu and Nakhleh, 2015b).

### Penalizing Network Complexity: Bayesian Inference

Discussing statistical inference in general, Attias (Attias, 1999) listed three problems with maximum likelihood: “First, it produces a model that overfits the data and subsequently have [sic] suboptimal generalization performance. Second, it cannot be used to learn the structure of the graph, since more complicated graphs assign a higher likelihood to the data. Third, it is computationally tractable only for a small class of models.” When the model of interest in a phylogenetic tree, the first two problems are generally not of concern (barring the complexity of the model of evolution underlying the inference). Putting aside the problem of computational tractability, the first two problems listed by Attias are an Achilles heel for phylogenetic network inference by maximum likelihood.

Phylogenetic networks can be viewed as mixture models whose components are distributions defined by parental trees of the network (Zhu et al., 2016). Inferring the true model in the case of phylogenetic networks includes determining, in addition to many other parameters, the true number of reticulations. Inference of such a model based on an unpenalized likelihood can rarely work since, as Attias pointed out, “more complicated graphs assign a higher likelihood.” In fact, the notion of *statistical consistency*, which has been a staple in the literature on phylogenetic tree inference, is not even applicable in the case of maximum (unpenalized) likelihood of phylogenetic networks—it is easy to imagine scenarios where adding more reticulations to the true phylogeny would only improve the likelihood. Therefore, in analyzing data sets under the aforementioned likelihood-based inference methods we recommend experimenting with varying the maximum allowed number of reticulation events and inspecting the likelihoods of the models with different numbers of reticulations.

A more principled way to deal with model complexity is via Bayesian inference, which allows, among other things, for regularization via the prior distribution. The **MCMC_GT** command performs Bayesian inference of the posterior distribution of the species network along with its branch lengths (in coalescent units) and inheritance probabilities via reversible-jump Markov chain Monte Carlo (RJMCMC). The input consists of single or multiple rooted gene tree topologies per locus, as above, and the number of individuals per species could vary across loci. Full details of the Bayesian inference of phylogenetic networks can be found in (Wen et al., 2016).

To handle gene tree uncertainty in a principled manner, and to allow for inferring the values of various network-associated parameters, PhyloNet implements Bayesian inference of phylogenetic networks directly from sequence data. The **MCMC_SEQ** command performs Bayesian inference of the posterior distribution of the species network along with its divergence times (in units of expected number of mutations per site) and population mutation rates (in units of population mutation per site), inheritance probabilities, and ultrametric gene trees along with its coalescent times (in units of expected number of mutations per site) simultaneously via RJMCMC. The input consists of sequence alignments of unlinked loci. Multiple individuals per species could be used, and their numbers could vary across loci. Full details of the co-estimation method can be found in (Wen and Nakhleh, 2017). The **MCMC_BiMarkers** command, on the other hand, performs Bayesian inference of the posterior distribution of the species network along with its divergence times (in units of expected number of mutations per site), population mutation rates (in units of population mutation per site), and inheritance probabilities via RJMCMC. The input consists of bi-allelic markers of unlinked loci, most notably single nucleotide polymorphisms (SNPs) and amplified fragment length polymorphisms (AFLP). Also, multiple individuals per species could be used. This method carries out numerical integration over all gene trees, which allows it to completely sidestep the issue of sampling gene trees. Full details of the computation can be found in (Zhu et al., 2017).

### Other Features

In addition to the aforementioned inference methods and all the functionalities that existed in the old version of PhyloNet (Than et al., 2008), the software package includes new features that help with other types of analyses.

As trees are a special case of networks, all the features above allow for species tree inference by simply setting the number of reticulations allowed during the analysis to 0. Additionally, PhyloNet implements greedy consensus, the “democratic vote,” and the GLASS method of (Mossel and Roch, 2010).

PhyloNet also includes a method for distance-based inference of phylogenetic networks (Yu and Nakhleh, 2015a), as well as a Gibbs sampling method for estimating the parameters of a given phylogenetic network (Yu et al., 2016).

Last but not least, the **SimGTinNetwork** and **SimBiMarkersinNetwork** simulate gene trees and bi-allelic markers, respectively, on a phylogenetic network. In particular, the former automates the process of simulating gene trees in the presence of reticulation and incomplete lineage sorting using the program ms (Hudson, 2002). The latter command extends the simulator developed in (Bryant et al., 2012).

## INPUT AND OUTPUT FORMATS

PhyloNet 3 is a software package in the JAR format that can be installed and executed on any system with the Java Platform (Version 7.0 or higher). The command line in a command prompt is

~~~
java -jar PhyloNet_X.Y.Z.jar script.nex
~~~

where **X.Y.Z** is the version number (version 3.6.1 is the most recent release), and **script.nex** is the input NEXUS file containing data and the PhyloNet commands to be executed.

The input data and the commands are listed in blocks. Each block start with the “BEGIN” keyword and terminate with the “END;” keyword. Commands in a PHYLONET block begin with a command identifier and terminate with a semicolon.

In the example input file below, to estimate the posterior distribution of the species network and the gene trees, the input sequence alignments are listed in the “DATA” block. The starting gene tree for each locus and the starting network, which is optional, can be specified in the “TREES” block and the “NETWORKS” block, respectively. Finally the command *MCMC_SEQ* and its parameters are provided for execution in the “PHYLONET” block. Details about specific parameters for a given command can be found on the website of PhyloNet.

~~~
#NEXUS

BEGIN DATA;
    Dimensions ntax=3 nchar=35;
    Format datatype=dna symbols=“ACTG” missing=?
     gap=−;
    Matrix
[locus0, 15]
A TCGCGCTAACGTCGA
B GCGCACCTACTGCGG
B GCGCACCTACTCCGG
[locus1, 20]
A GAAACGGATCTAAGTGTACG
B CGCTCGGATCTAAGTGTACG
C CGCTCGGATCTAAGTGTACG
;END;

BEGIN TREES;
Tree gt0 = (A:0.119,(B:0.058,C:0.058):0.061);
Tree gt1 = (A:0.068,(B:0.016,C:0.016):0.052);
END;

BEGIN NETWORKS;
Network net = (((B:0.0)#H1:0.05::0.8,(C:0.002,
  #H1:0.002::0.2):0.048):0.01,A:0.06);
END;

BEGIN PHYLONET;
MCMC_SEQ -sgt (gt0,gt1) -snet net1 -sps 0.04;
END;
~~~

Gene trees are given in the Newick format, where the values after the colons are the branch lengths. Networks are given in the Rich Newick format which contains hybridization nodes denoted in “#H1”, “#H2”, …, “#Hn”, where *n* is the number of reticulations in the network. The branch lengths, the population mutation rates, and the inheritance probabilities are specified after the first, second and third colons, respectively. Note that the units of branch lengths (for both trees and networks) can either be coalescent units, or the number of expected mutations per site, depending on the requirement of PhyloNet command. The population mutation rate is optional if the branch lengths are in the coalescent units, or a constant population mutation rate across all the branches is assumed. The inheritance probabilities are only relevant for the hybridization nodes.

The **InferNetwork_MP** command returns species networks and the corresponding extra lineages. The **InferNetwork_ML** command and its relatives return species networks and the corresponding likelihood values. The total number of returned networks can be specified via *-n* option for both commands. As we stated above, the user can either specify the maximum number of reticulation events or utilize the cross-validation, bootstrap, information criteria to determine the model complexity when using maximum likelihood approach.

For Bayesian inference, the program outputs the log posterior probability, likelihood and prior for every sample. When the MCMC chain ends, the overall acceptance rates of the RJMCMC proposals and the 95% credible set of species networks (the smallest set of all topologies that accounts for 95% of the posterior probability) are reported. For every topology in the 95% credible set, the proportion of the topology being sampled, the maximum posterior value (MAP) and the corresponding MAP topology, the average posterior value and the averaged (branch lengths and inheritance probabilities) network are given. The model complexity is controlled mainly by the Poisson prior on the number of reticulations (see (Wen et al., 2016) for details). The Poisson distribution parameter can be tuned via “-pp” option.

## CONCLUSION

PhyloNet 3 is a comprehensive software package for phylogenetic network inference, particularly in the presence of incomplete lineage sorting. It implements maximum parsimony, maximum likelihood, and Bayesian inferences, in addition to a host of other features for analyzing phylogenetic networks and simulating data on them. The package is implemented in Java and is publicly available as an executable as well as source files.

In terms of the main aim of PhyloNet, which is the inference of phylogenetic networks, very few tools exist. TreeMix (Pickrell and Pritchard, 2012) is a very popular tool in the population genetics community. It uses allele frequency data and mainly targets analyses of admixtures among sub-populations of a single species. More recently, Bayesian inference of phylogenetic networks was implemented in BEAST2 (Zhang et al., 2017) and inference of unrooted networks based on pseudo-likelihood was implemented in the PhyloNetworks software package (Solís-Lemus et al., 2017). However, in terms of implementing inference under different criteria and from different types of data, PhyloNet 3 is the most comprehensive.

As highlighted above, the challenge of computational tractability aside, the major challenge with network inference in general is determining the true number of reticulations and guarding against overfitting. In particular, phylogenetic networks are more complex models than trees and can always fit the data at least as well as trees do. Does this mean networks are simply over-parameterized models that should be abandoned in favor of trees? We argue that the answer is no. First, if the evolutionary history is reticulate, a tree-based method is unlikely to uncover the reticulation events (their number and locations). Second, even if one is interested in the species tree “despite reticulation,” a species tree inference method might not correctly recover the species tree (Solís-Lemus et al., 2016). Third, even when the true evolutionary history is strictly treelike, the network structure could be viewed as a graphical representation of the variance around the tree structure. Insisting on a sparse network for convenience or ease of visual inspection is akin to insisting on a well-supported model no matter what the data says. Needless to say, one is interested in the true graph structure and not one that is more complicated simply because it assigns higher likelihood to the data. From our experience, the Bayesian approaches handle this challenge very well.

While we continue to improve the features and user-friendliness of the software package, the main direction we are currently pursuing is achieving scalability of the various inference methods in PhyloNet 3 to larger data sets in terms of the numbers of taxa as well as loci.

## AVAILABILITY

PhyloNet is publicly available for download from https://bioinfocs.rice.edu/phylonet. It can be installed and executed on any system with the Java Platform (Version 7.0 or higher). The current release includes source code, tutorials, example scripts, list of commands and useful links.

## FUNDING

Development of the various functionalities in PhyloNet has been generously supported by DOE (DE-FG02-06ER25734), NIH (R01LM009494), and NSF (CCF-0622037, DBI-1062463, CCF-1302179, CCF-1514177) grants and Sloan and Guggenheim fellowships to L.N. Author contributions: D.W., Y.Y., and J.Z. did almost all the implementation and testing of the inference features in PhyloNet 3; D.W., J.Z. and L.N. wrote the paper.

## ACKNOWLEDGEMENTS

We would like to thank the PhyloNet users for submitting bug reports, feature requests, and other comments that helped us significantly improve the software.

